# Preferred walking speed on rough terrain; is it all about energetics?

**DOI:** 10.1101/332106

**Authors:** Koren Gast, Rodger Kram, Raziel Riemer

**Author notes:** Corresponding author: Raziel Riemer.

## Abstract

Humans have evolved the ability to walk very efficiently. Further, humans prefer to walk at speeds that approximately minimize their metabolic energy expenditure per unit distance (i.e. gross cost of transport, COT). This has been found in a variety of population groups and other species. However, these studies were performed on smooth, level ground or on treadmills. We hypothesized that the objective function for walking is more complex than only minimizing the COT. To test this idea, we compared the preferred speeds and the relationships between COT and speed for people walking on both a smooth, level floor and a rough, natural terrain trail. Rough terrain presumably introduces other factors, such as stability, to the objective function. 10 healthy men walked on both a straight, flat, smooth floor and on an outdoor trail strewn with rocks and boulders. In both locations, subjects performed 6-8 trials at different speeds relative to their preferred speed. The COT-speed relationships were similarly U-shaped for both surfaces, but the COT values on rough terrain were approximately 215% greater. On the smooth surface, the preferred speed (1.24+/-0.14 m/sec) was 5.1% faster (p>0.05) than the speed that minimized COT (1.18 m/sec) but on rough terrain, the preferred speed (1.06+/-0.07 m/sec) was 6.2% slower (p>0.05) than the COT minimum speed (1.13 m/sec). This suggests that the objective function when walking on rough terrain includes additional factors such as stability.

Summary statement: On rough terrain, humans do not choose their walking speed based on metabolic energy minimization alone.

## Introduction

Humans have evolved the ability to walk very efficiently. Over generations, our bodies have evolved muscular and skeletal systems well-suited to locomotion (Alexander, 2003). Further, we learn and choose to walk in a way that minimizes our metabolic energy expenditure (Ralston, 1958; Zarrugh et al., 1974). For example, it has been shown that step frequency (Zarrugh et al., 1974), step length (Umberger and Martin, 2007), step width (Donelan et al., 2001) and speed (Zarrugh et al., 1974) are all chosen to minimize energy expenditure.

More specifically, it has been found that humans choose a walking speed (i.e. preferred speed) that is close to the metabolically optimal speed which minimizes the Cost of Transport (COT) - the metabolic rate divided by the locomotion speed. This phenomenon has been observed in people of normal weight, in people who are with obese (Browning et al., 2006), in people with trans-tibial and trans-femoral amputations (Genin et al., 2005), in people with post-polio syndrome (Ghosh et al., 1982), and when people carry loads (Bastien et al., 2005). These studies all support the idea that while walking, our body optimizes *MIN*_{θ}_[*COT(*θ)], where θ is a vector of walking parameters.

However, there are exceptions to this rule. For example, Clark-Carter et al. (1986) found that people who are blind prefer walking speeds similar to sighted people when they are accompanied by a guide. But without a guide, their prefer walking speed was slower and likely not at their COT minimum. More recently, it has recently been discovered that when walking downhill, humans do not select a gait pattern that minimizes COT. Monsch et al. (2012) found that when instructed to walk in a “loose relaxed gait” subjects had a lower COT than when walking with their natural, preferred gait. Similarly, it was found that people walk more slowly on a smooth surface when it is elevated to above the ground (Brown et al., 2002; Schnjepp et al., 2014) and thus presumably they chose to not walk at the energetic COT minimum.

These studies led us to propose that humans choose walking parameters to optimize an objective function that is more complex than *MIN*_{θ}_[*COT(*θ)]. For example, this objective function could take the form:

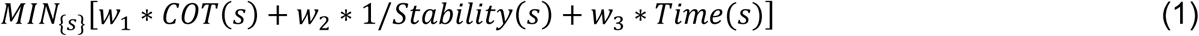

where s is the walking speed, and w_1_, w_2_ and w_3_ are weighting coefficients that represent the importance of the different factors for a given task. This formulation proposes that when choosing walking parameters, we optimize not only for COT but also for stability and time of completion. This idea is in agreement with Shadmehr et al. (2016) who proposed an objective function for humans while performing a reaching motion is different from COT minimization alone.

So far, most research into walking and the COT phenomenon has been carried out on treadmills or smooth, level floors. However, there are several studies that investigate the metabolic rate of locomotion on natural terrains but without the focus on optimal speed and preferred speed. For example, walking has been investigated on sand (Pinnington and Dawson, 2001), grass (Davies and Mackinnon, 2006), dirt roads (Daniels et al., 1953) and snow (Pandolf et al., 1976; Soule and Goldman 1972). Givoni and Goldman, (1971), and Pandolf et al. (1977) developed predication equations for the metabolic cost of load carrying while walking on different terrains and slopes. A recent study examined the metabolic cost of walking on a treadmill that imitates uneven terrain (Voloshina and Ferris, 2013). However, to the best of our knowledge, no one has examined the COT as a function of speed on natural, rough terrain.

In this study, we compared the COT during walking at different speeds on a smooth level floor and over natural, rough terrain. Investigation of COT on natural surfaces is important for two main reasons. First, human walking efficiency primarily evolved on natural surfaces, not smooth floors or treadmills. Second, walking on rough terrain intrinsically requires the person to consider their stability while walking. Therefore, we hypothesized that on rough terrain, the preferred walking speed would be slower compared to smooth and the preferred speed would be slower than the metabolic COT minimum speed. If this is found to be the case, it would imply that the objective function for human walking does include some sort of “stability” factor, which is greater on rough terrain than on smooth level surfaces.

## Methods

### Subjects

10 healthy male students (body mass: 76.5 ± 11.5 kg, age: 27.5 ± 1.6 y; mean ± 1 S.D.) participated in this experiment. All test subjects were instructed to sleep for at least six hours on the night prior to the experiment and to eat a light breakfast ending at least two hours prior to the start. The Ben-Gurion University IRB committee approved the study and participants gave informed consent. Each subject performed three walking sessions – one session on a smooth, level floor and two sessions on rough terrain.

### Protocol

For the smooth level concrete floor condition, the route was 44m long, straight, flat, and in the shade. For the rough terrain condition, the subjects walked out-and-back along a 67m long trail, measured along the path itself, not point-to-point. The path was relatively straight and it undulated with a maximum elevation amplitude of less than 2 m. The trail comprised some randomly scattered rocks and boulders (**Fig. 1**).

**Fig. 1.**
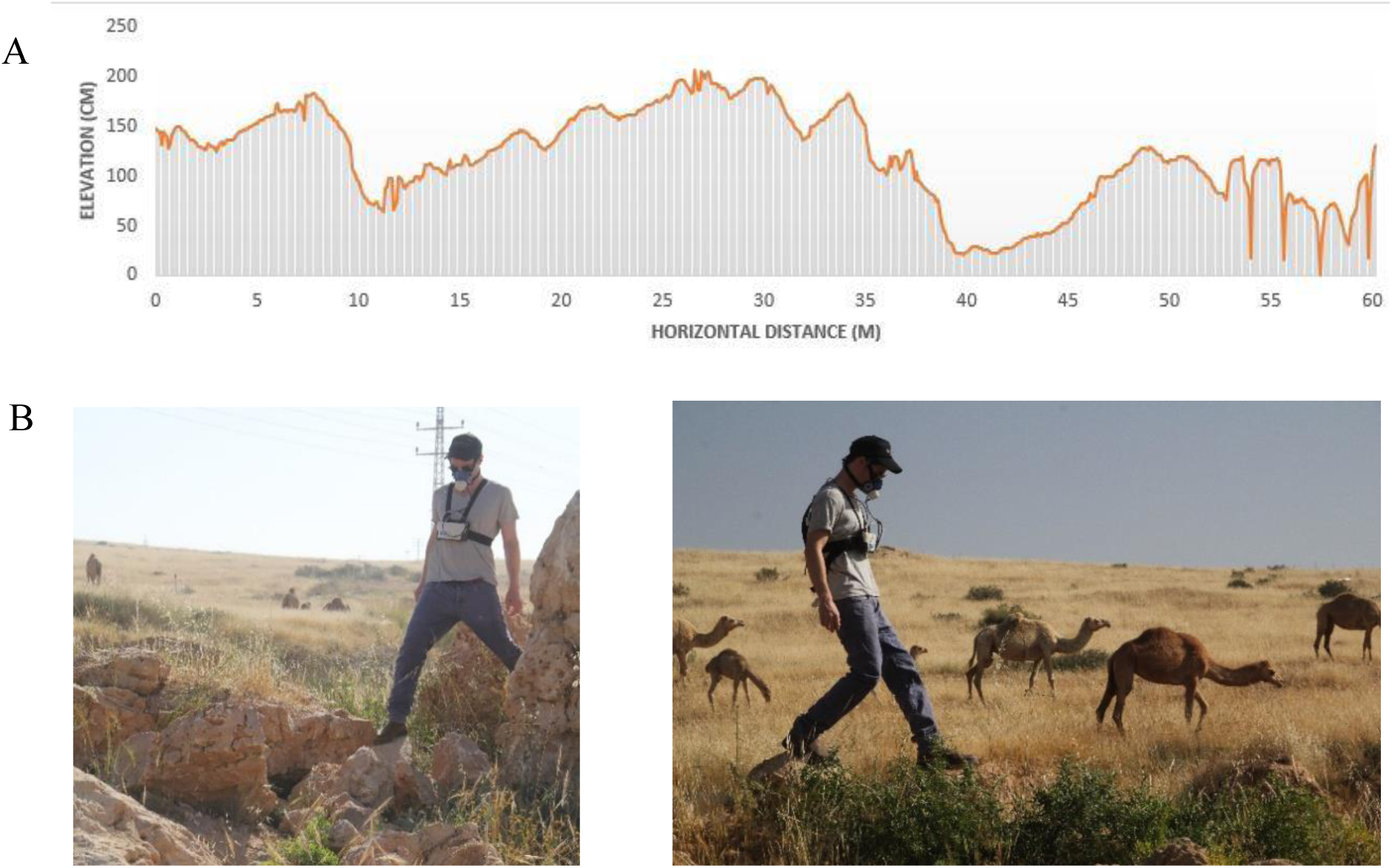
The rough terrain path. Side view of the trail (A) showing the elevation profile at 10 cm horizontal intervals. These are relative elevation values, where the lowest point is assigned an elevation of 0. Subjects sometimes stepped over deep, narrow cracks or small, sharp rocks. B. Pictures of one of the subjects walking on the trail.

### Experimental procedure

Subjects did not need any practice to walk comfortably on the smooth floor. However, the rough terrain condition required practice. Although all of the subjects had prior experience of rough terrain walking, they do not partake in this activity daily. In a pilot study, we found that there was a learning effect and that it took approximately 25 min of practice before the COT stabilized. Therefore, to eliminate the possibility of acquiring data during this learning period, subjects completed two rough terrain sessions in which the subject performed the full experimental protocol. For our analysis, we used only the second session. At the beginning of each session, after the subject had been fitted with the metabolic measurement system, they walked out-and-back along the trail with a guide (one of the research team members) who showed them the route which was marked with small flags. Subjects then walked the trail by themselves for at least 10 min at various speeds, starting with their preferred speed, followed by 10 min at their maximum walking speed and then finally 10 min at a very slow speed (about 50% of their preferred speed). At each speed, the subjects completed at least one full out-and-back circuit. In a pilot study, we found that this protocol accelerated learning and reduced adaptation time. The protocol was inspired by Selinger et al. (2015) who studied humans walking with novel exoskeletons and found that in order to find the optimal step frequency that minimized their metabolic rate, subjects had to carry out an exploratory session in which they walked at fast and slow step frequencies.

After the practice trial, the main experiment started. First, the subjects walked on the trail for 7-9 minutes at their preferred speed. We calculated the average speed (i.e. preferred speed) from the time for completion of the trail’s known distance. Next, the subjects performed a total of 6-8 trials at approximately 50, 75, 100 and 150% of their preferred speed. 150% of preferred speed was close to their maximum walking speed. Thus, some of the speeds were performed more than once. There was a 5-minute rest period between trials. Walking speed was coached via verbal commands from the researcher. We randomized the order of speeds for each subject to avoid trial order effects. In a pilot study, we found that steady-state rates of energy expenditure were obtained after about 1.5-2 minutes. Therefore, we only analyzed measurements of metabolic rate that were obtained at least 3 minutes after the start time. To eliminate the effects of local terrain variation, we averaged metabolic rate values over full out-and-back laps. The same procedures were employed for the smooth floor condition.

We measured the metabolic energy consumption using a K4b2 telemetric indirect calorimetry system (Cosmed, Rome, Italy). This system includes a portable unit which consists of a processing unit containing the O_2_ and CO_2_ analyzer and a battery pack. Together, the unit has a mass of 1.5 kg and was worn by the subject along with a silicone mask containing a flow-rate turbine. Every day and before each trial, the turbine was calibrated using a standard authorized calibration gas mixture and a volume pump.

### Data analysis

Metabolic rate was calculated using the Brockway (1987) equation. We then calculated the COT by dividing the average metabolic rate by the average speed and the subject’s body mass, i.e.

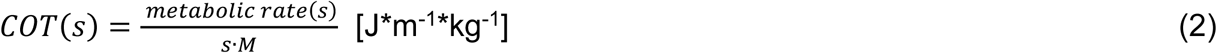

where *s* is the walking speed and M is body mass. To ensure that the metabolic energy was primarily generated via aerobic metabolism, only trials with an RER (respiratory exchange ratio) of less than 1.00 were analyzed.

To derive a model to predict the metabolic rate as a function of walking speed, we tested polynomials of orders 1 to 4 to find the best fit to the data (highest R^2^) for each surface’s dataset. To test whether there was a significant difference in the relationship between metabolic rate and speed between the two surfaces, we applied a Chow test to determine whether if there was a difference between the regression equations of the COT on both surfaces:

The Chow test F statistic was calculated as:

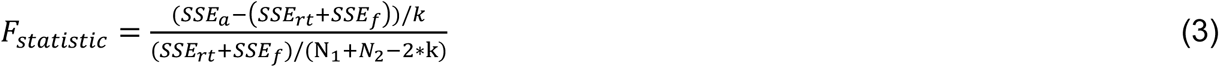

where SSE_a_ is the SSE (sum squared of errors) calculated on all the observations, SSE_rt_ is the SSE of the observations from the rough terrain experiments, SSE_f_ is the SSE of the observations from the experiments on the floor surface, k = 3 is the number of variables in the regression equations and N_1_ = 61 (rough terrain), N2 = 57 (floor) are the number of observations in each group.

To test whether there was a significant difference between the preferred and energetically optimal speeds for each surface, we used a paired t-test to compare 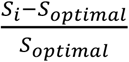 in the floor condition to the same index in the rough terrain condition, where s_i_ is the preferred walking speed of subject i and s_optimal_ is the speed at the minimal COT value calculated using the fitted function for each surface (the speed where the function’s first derivative equals zero).

Finally, we used another Chow test to determine whether the two terrains had differently shaped COT vs. speed relationships. To perform this comparison, all values were normalized as follows. For each subject, all walking speeds were divided by the subject’s preferred speed and all COT values were divided by the COT of the subject at his preferred walking speed.

## Results

### Metabolic rate

After testing regression models with 4 different polynomials orders, we found that for walking on both the smooth floor and rough terrain conditions, the best fit to the data (i.e. largest R^2^) was in the form: Metabolic rate(s) = as^2^ + bs + c where metabolic rate is normalized to mass (W/kg) and s is the walking speed

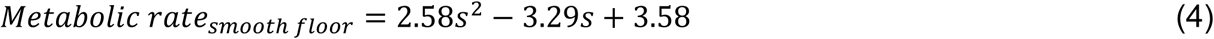

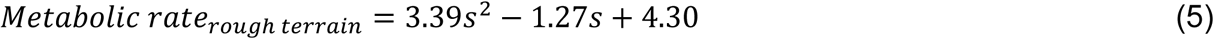

For Equation 4, the R^2^ value was 0.92 and for Equation 5 the value was 0.85. The quadratic form of the equation for predicting metabolic rate as a function of speed is in agreement with the results of Ralston (1958) and Zarrugh, Todd, & Ralston (1974). For a linear increase in walking speed, we observe a polynomial increase in metabolic rate (**Fig. 2**). The Chow-test showed that there was a significant difference between the smooth and rough terrain conditions (p-value<10^−15^); therefore it was reasonable to fit a different set of function constants for each surface.

**Fig. 2.**
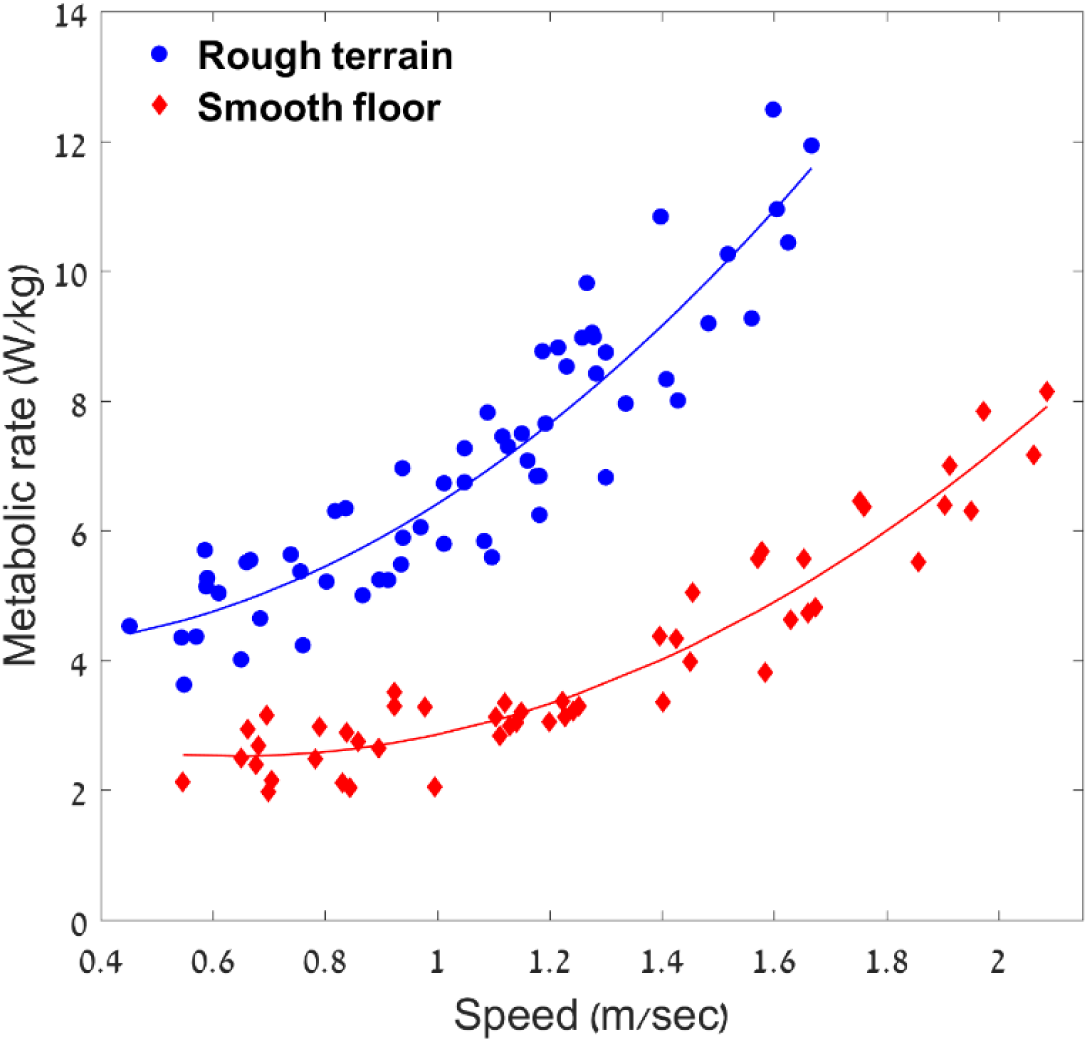
Metabolic rate as a function of walking speed. Data of total 118 trials on the rough terrain (61 trials) and floor (57 trials) surfaces, obtained from 10 subjects walking at a variety of speeds.

### COT

To calculate the COT functions, we divided the metabolic rate functions (Equations 4, 5) by walking speed (s):

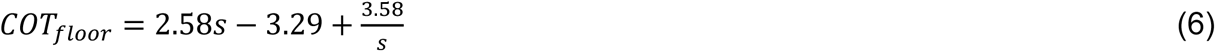

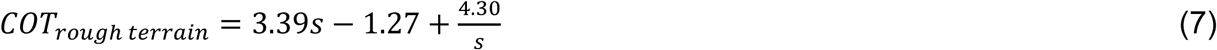

We found that the average COT values for the rough terrain were approximately 215% greater than those obtained for the level floor condition. The preferred walking speed (averaged across all subjects) on rough terrain was about 85% of the preferred walking speed on the smooth floor (**Table 1 and Fig. 3**).

**Table 1:** Metabolic results as a function of walking speed and terrain (mean ± 1 s.d.)

| speed category: |  | 50% | 75% | 100%* | 125% | max |
| --- | --- | --- | --- | --- | --- | --- |
| Smooth floor | Speed ($\frac{m}{sec}$) | 0.68±0.06 | 0.87±0.05 | 1.24±0.14 | 1.6±0.12 | 1.98±0.08 |
| | Metabolic rate<br>($\frac{Watt}{kg}$) | 2.56±0.38 | 2.79±0.47 | 3.46±0.83 | 4.77±1.01 | 7.06±0.89 |
| | COT ($\frac{J}{kg*m}$) | 3.78±0.51 | 3.19±0.42 | 2.75±0.37 | 2.95±0.43 | 3.56±0.34 |
| Rough terrain | Speed ($\frac{m}{sec}$) | 0.57±0.05 | 0.78±0.07 | 1.07±0.07 | 1.3±0.07 | 1.52±0.12 |
| | Metabolic rate<br>($\frac{Watt}{kg}$) | 4.67±0.62 | 5.44±0.57 | 6.61±0.73 | 8.41±0.65 | 10.58±1.05 |
| | COT ($\frac{J}{kg*m}$) | 8.24±1.22 | 7.04±1.00 | 6.21±0.65 | 6.51±0.6 | 6.96±0.67 |

\*100% is the preferred speed

**Fig. 3.**
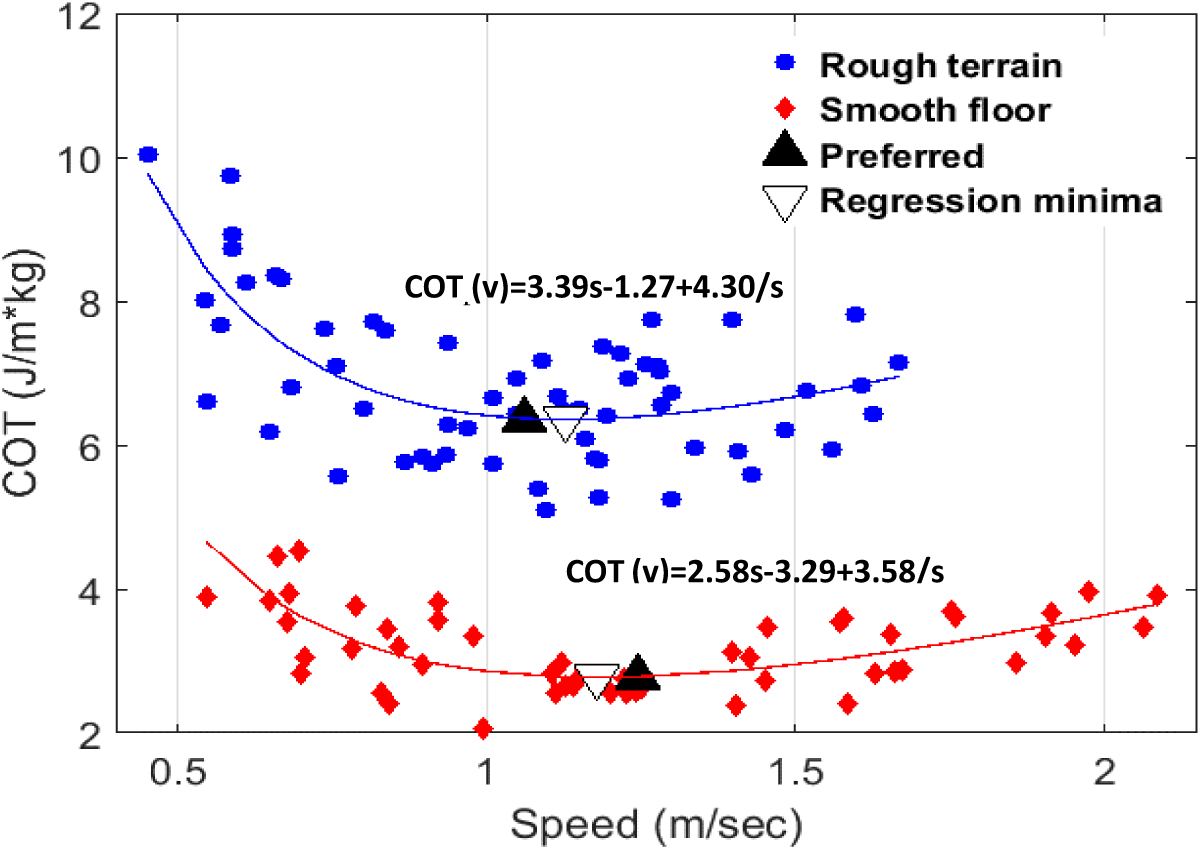
COT as a function of walking speed for all trials.

The main goal of this study was to compare how humans choose their preferred walking speed on smooth and rough terrain. The average preferred speed was 1.24 m/sec (+/-0.14) on smooth terrain and 1.06 m/sec (+-0.07) on the rough terrain. The metabolically optimal speeds, the speed that minimizes the fitted COT functions, were 1.18 m/sec on the smooth floor and 1.13 m/sec on rough terrain. The preferred speed was close to the optimum speed for both surfaces. However, the preferred speed was faster than the optimum speed for the smooth floor condition (by 5.1%), while on rough terrain, the preferred speed was slower than the optimum (−6.2%). A paired t-test revealed that these speed differences (i.e. the gap between preferred speed and optimal speed on floor compared to on rough terrain) were statistically significant (p-value<0.05).

### COT change as a function of speed

We tested whether the COT vs. speed functions differed between the terrain conditions. After normalizing the COT and speed values, we derived equations 8 and 9:

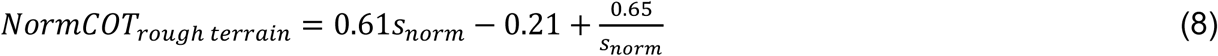

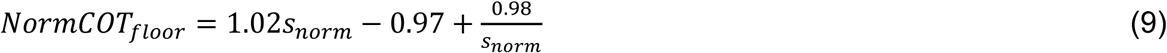

where NormCOT are the COT values of a subject divided by the COT at their preferred speed and *s*_*norm*_ is the walking speed of a subject divided by their preferred walking speed. The normalized COT data are shown in **Fig. 4**. The Chow test did not show a significant difference (p=0.28) between the shape of the two curves.

**Fig. 4.**
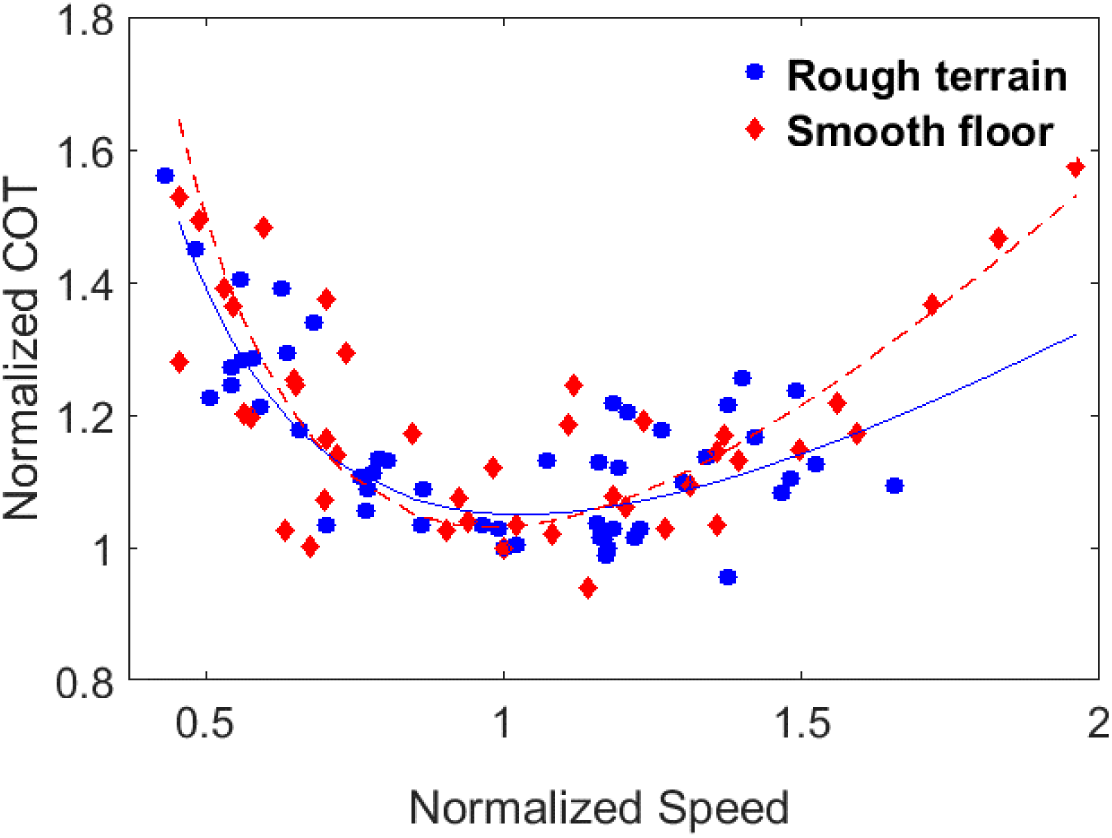
Comparison of normalized COT functions for the smooth floor and rough terrain conditions.

## Discussion

The aim of this study was to compare the preferred speeds and energetic COT for humans walking on smooth and rough terrain across a range of walking speeds. The cost of transport for the rough terrain was considerably greater (∼215%) than the smooth level walking, a finding that is very similar to past research for walking on sand dunes (Givoni and Goldman, 1971). The differences in COT values for different surfaces likely reflect: different rates of mechanical work performed on the center of mass and the substrate itself (Lejeune et al., 1998), greater decelerations and accelerations of the center of mass because of foot placement (Seethapathi and Srinivasan, 2015), changes in step length and step width (Donelan et al., 2001) and/or other stability-related issues. Voloshina and Ferris (2013) have shown that walking on uneven terrain causes only minor changes in stepping strategy and suggest that the changes in metabolic rate are instead due to a change in the amount of work carried out by lower-limb joints as well as changes in the timing of foot-ground collision and trailing leg push-off.

The function describing the dependence of COT on speed for the smooth level floor condition in the present study is similar to predication equations developed in previous studies (Givoni and Goldman, 1971; Ralston, 1958; Zarrugh et al., 1974; Pandolf et al., 1977). Our data for walking on the smooth floor most closely match the prediction equation of Givoni and Goldman (1971) (**Fig. 5A.**). For the rough terrain (**Fig. 5B.**), we compared our data to two prediction equations for metabolic rate from previous studies: Givoni and Goldman (1971) and Pandolf et al. (1977). The terrain factors (η) that resulted in the best match to our data and that were used to generate the curves in **Fig. 5B.** were 2.2 and 2.8 for Givoni and Goldman (1971) and Pandolf et al. (1977), respectively. The prediction equation of Givoni and Goldman (1971) was also the closest match to our rough terrain data. Although the Pandolf et al. (1977) equations are used more commonly, our data better matched the Givoni and Goldman (1971) equations. A likely reason is that Pandolf et al. (1977) developed their prediction equations based on walking speeds ranging only from 0.2 to 1.0 m/sec, while Givoni and Goldman (1971) investigated speeds from 0.55 to 2.50 m/sec which is similar to the speeds used in our experiment (0.45 to 2.20 m/s).

**Fig. 5.**
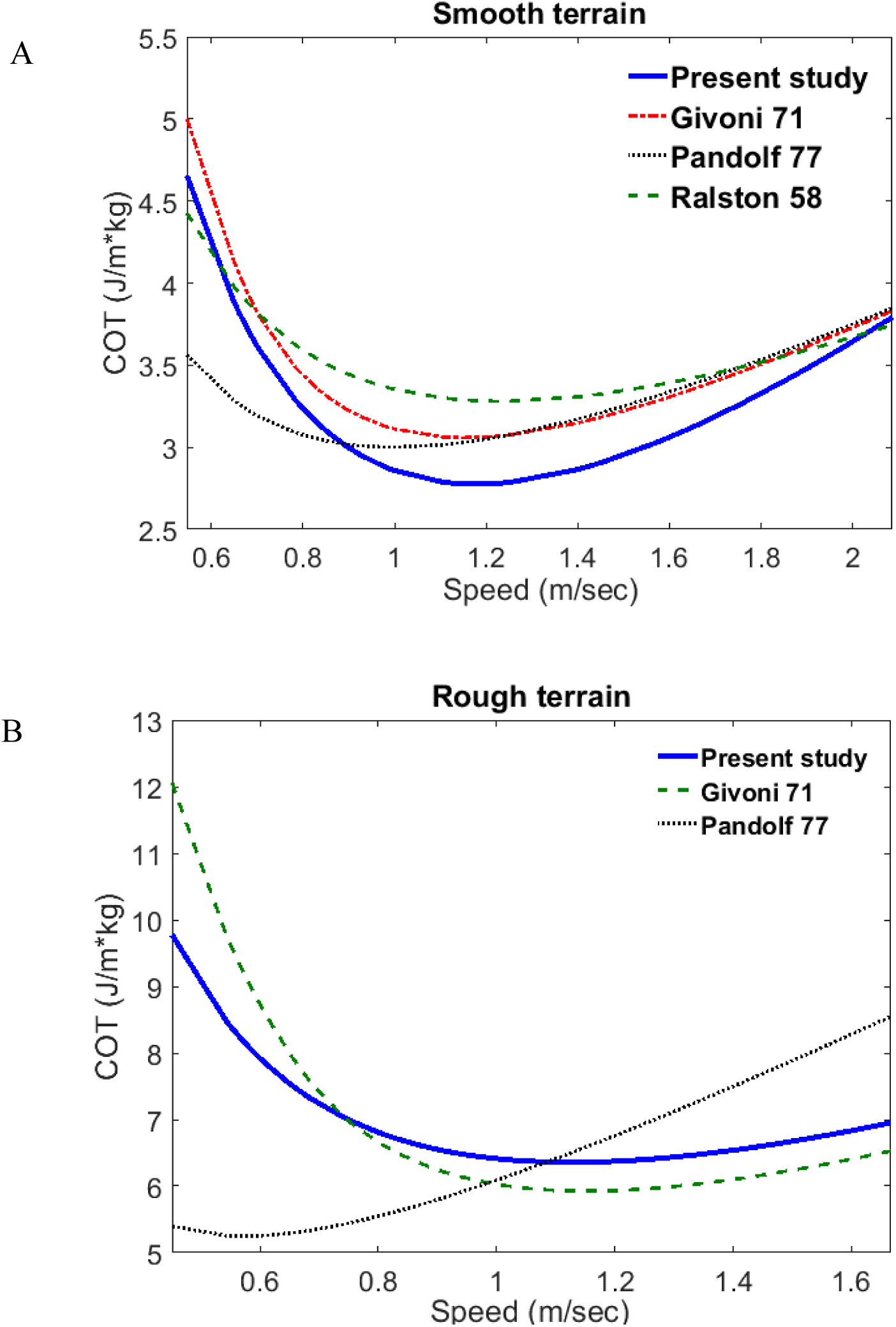
Comparison of our COT predication function with those reported previously. (A) Fitted curves of both past studies (Ralston, 1958; Givoni and Goldman, 1971; Pandolf et al., 1977) and the present study for COT values of walking on smooth surfaces. (B) Fitted curves to COT values from rough terrain (present study) and other surfaces (Givoni and Goldman, 1971; Pandolf et al., 1977). More of previous studies curves in **appendix 1**.

We hypothesized that the objective function that humans try to minimize is more complex than just COT and proposed Equation. 1 as one possible form. Specifically, since there was likely to be a difference in stability between the two conditions, we hypothesized that the relative preferred speed (preferred speed / optimal speed) on rough terrain would be slower than the relative preferred speed on smooth, level ground. Our results revealed that similar to past studies that studied COT on smooth, level ground and treadmills (e.g. Ralston, 1958; Zarrugh et al., 1974), the subjects’ preferred speeds on both surfaces were close to the speeds that minimized the COT. However, the preferred speed on the smooth floor was faster than the optimum speed (+5.1%), while on rough terrain, the preferred speed was slower than the optimum (−6.2%) with p-value<0.05. This finding supports our hypothesis that humans are probably optimizing a more complex function than *MIN*_{θ}_[*COT(*θ)]. To explain our results, we refer to Equation. 1; yet we do not claim that this is the only function possible. Based on this equation, we argue that when walking on rough terrain, the slower walking speed increases stability, thus reduces the overall value of the objective function. On the smooth, level floor, stability likely plays a smaller role and the time for completion of the task (the third factor in Equation. 1) is likely to play a larger role; therefore, subjects walk slightly faster than the speed corresponding to the minimum COT.

It is worth noting that other species, for example wildebeest (Pennycuick, 1975), elephants (Langman et al., 1995) and horses (Hoyt & Taylor, 1981), also exhibit preferred terrestrial locomotion speeds within each gait. Further, the preferred walking speeds of horses and elephants also are close to the minimum COT speeds (Hoyt & Taylor, 1981; Langman et al., 1995). It remains to be tested if factors other than COT affect the preferred speeds in these and other species.

The magnitude of the difference between the preferred speed and the energetically optimal COT speed for the two surfaces averaged 5.7%. However, the COT functions in **Fig. 3** are very shallow near the optimum speed such that a difference of 5.7% in speed results in a change in COT value of only ∼0.2%. It is worth pondering if and how humans can accurately sense such a small difference in COT. After all, COT requires information about instantaneous metabolic rate and walking speed. Humans can reliably perceive their physiological effort, presumably via cardiac and pulmonary sensors (Borg, 1982) or localized effort which can be reflected in electromyographic recordings (Korol at el 2017). More specific to locomotor optimization, Wong et al., 2017 investigated whether people utilize the body’s blood gas receptors to identify their optimal step frequency. The investigators experimentally manipulated blood gas (O_2_ and CO_2_) concentrations and found that their subjects ignored the blood gas receptor information and walked with their normal step frequencies. Mohler et al., (2007) has shown that human perception of and selection of preferred walking speed is influenced by vision. Thus, it seems possible that humans prefer certain walking speeds based on instantaneous sensations of effort and speed and thus minimum COT, but it is also possible that past walking experience sets the baseline walking speed.

In summary, based on both the current findings and previous research (Brown et al., 2002; Clark-Carter et al. 1986; Monsch et al., 2012; Schnjepp et al., 2014).

It is seems that simply minimizing the Cost of Transport does not fully represent the human objective function in walking. Other walking conditions should be examined to investigate additional parameters that might appear in the cost function, such as stability, reward and time-saving.

List of Symbols and Abbreviations

| Symbol | Meaning |
| --- | --- |
| COT | Cost of transport |
| NormCOT | Normalized cost of transport |
| SSE | Sum squared of errors |
| s | speed |
| $s_{norm}$ | Normalized speed |

## Acknowledgements

We thank Prof. Yisrael Parmet for his help with the statistics for this study and Yosef Zahavi and Ori Wildikan for their help with the experiments.

## Author contributions

K.G., R.K. and R.R. conceived of the experiment. K.G. R.K. and R.R developed the experimental setup. K.G. and R.R. ran the pilot experiment and K.G. ran the main experiments. K.G. performed the statistical analyses. All the authors discussed the results and contributed to the final manuscript.

### Appendix 1: Details on how we generated previous studies COT regression curves

In Pandolf et al 1977 the metabolic rate fitted curve is in form of:

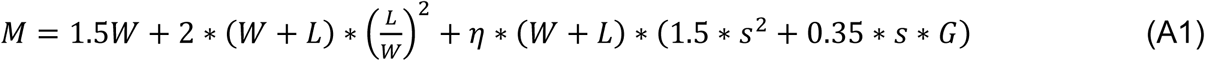

Where M is the metabolic rate in watt, W is the body weight (for this study the average body weight is 76.5 kg), L is external load which in our case is 1.5 kg for the metabolic rate measurement system (K4b2, Cosmed, Rome, Italy). G is the slope. In our case 0%. η is the terrain factor. η = 1 for smooth floor and η = 3 for rough terrain, this value was chosen to as it minimized the root mean square error (RMSE) between the experiment regression fit and Pandolf et al 1977 predication. S is the walking speed in m/s. After assigning the values to the equation A1 we obtained:

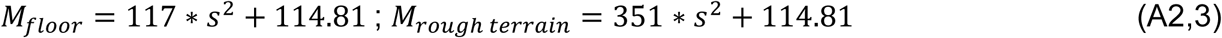

We divide M by (W+L) to obtain the metabolic rate per 1kg and by s to get the COT:

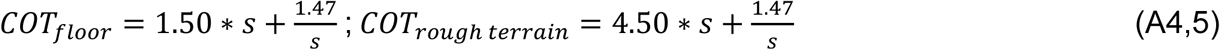

In Givoni and Goldman 1971 the metabolic rate regression curve is in form of:

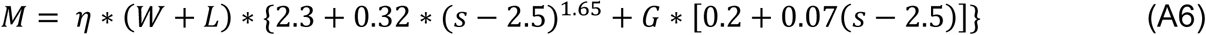

Where M is the metabolic rate in kcal/hr, W is the body weight (for this study the average body weight is 76.5 kg), L is external load which in our case is 1.5 kg for the metabolic rate measurement system (K4b2, Cosmed, Rome, Italy). G is the slope. In our case 0%. η is the terrain factor. η = 1 for smooth floor and η = 1.9 for rough terrain, chosen to minimize the RMSE. S is the walking speed in km/hr. After converting to the units used in the present study and assigning the above values to equation A6, we obtain:

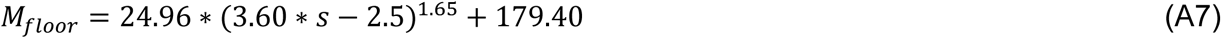

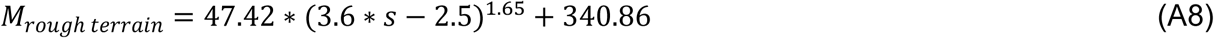

We divide M by (W+L) to obtain the metabolic rate per 1kg and by s to get the COT:

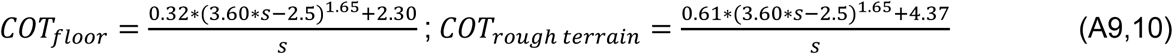

In Ralston 1958 the energy expenditure regression curve is in form of:

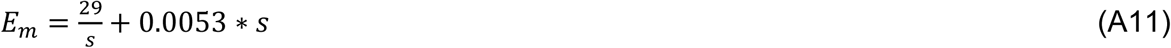

Where *E*_*m*_ is the energy expenditure in cal/meter/kg, and S is the walking speed in m/min which will be converted to m/s. After converting to the units used in the present study and assigning the above values to equation A11, we obtain:

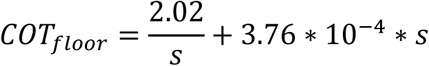

